# Morphological diversity of blastula formation and gastrulation in temnopleurid sea urchins

**DOI:** 10.1101/047472

**Authors:** Chisato Kitazawa, Tsubasa Fujii, Yuji Egusa, Miéko Komatsu, Akira Yamanaka

**Affiliations:** Biological Institute, Faculty of Education, Yamaguchi University, Yoshida 1677-1, Yamaguchi 753-8513, Japan; Biological Institute, Graduate School of Education, Yamaguchi University, Yoshida 1677-1, Yamaguchi 753-8513, Japan; Department of Biology, Graduate School of Science and Engineering for Research, University of Toyama, Toyama 930-8555, Japan; Laboratory of Environmental Biology, Graduate School of Medicine, Yamaguchi University, Yoshida 1677-1, Yamaguchi 753-8512, Japan

**Keywords:** primary mesenchyme cells ingression, extracellular matrix, blastular wall, cell morphology, gut elongation

## Abstract

Embryos of temnopleurid sea urchins exhibit species-specific morphologies. While *Temnopleurus toreumaticus* has a wrinkled blastula, others have a smooth blastula. Embryos of *T. toreumaticus* invaginate continuously at gastrulation, whereas in some others invagination is stepwise. We studied blastula and gastrula formation in four temnopleurids using light and scanning electron microscopy to clarify the mechanisms producing these differences. Unlike *T. toreumaticus*, blastomeres of mid-blastulae in *T. reevesii*, *T. hardwickii* and *Mespilia globulus* formed pseudopods. Before primary mesenchyme cells ingressed, embryos developed an area of orbicular cells in the vegetal plate. The cells surrounding the orbicular cells extended pseudopods toward the orbicular cell area in *T. toreumaticus*, *T. reevesii* and *T. hardwickii*. In *T. toreumaticus*, the extracellular matrix was well-developed and developed a hole-like structure that was not formed in others. Gastrulation of *T. reevesii*, *T. hardwickii* and *M. globulus* was stepwise, suggesting that differences of gastrulation are caused by all or some of factors: change of cell shape, rearrangement, pushing up and towing of cells. These species-specific morphologies may be caused by the shape and surface structure of blastomeres with cell-movement.

**Summary statement:** Temonopleurid embryology

## INTRODUCTION

Embryos of many sea urchins exhibit some species-specific morphological difference at the blastula and gastrula stages. Most indirect-developing species, which develop from a small egg with little yolk through planktotrophic larval stages, form a blastula with a smooth blastular wall, whereas direct-developing species, which develop from a large yolky egg either lack or undergo an accelerated larval stage and form a wrinkled blastula (Raff, 1987). These differences in formation are also known in other echinoderms (Henry et al., 1991).

Another important morphogenesis is gastrulation. After the primary mesenchyme cells (PMCs) are released into the blastocoel, the vegetal plate invaginates into the blastocoel to form the main internal structures including the archenteron (Trinkaus, 1984). There are least five steps of gastrulation in sea urchin embryos; formation of a thickened vegetal plate, primary invagination to form a gut rudiment, elongation of the gut rudiment and appearance of secondary mesenchyme cells (SMCs), secondary invagination to elongate more until reaching the internal surface of the apical plate, and tertiary invagination to recruit presumptive endodermal cells (Dan and Okazaki, 1956; Gustafson and Kinnander, 1956; Kominami and Takata, 2004). The manner of invagination of the archenteron in sea urchins is into two types: stepwise or continuous invagination (Kominami and Masui, 1996). Species of the stepwise type pass the first and secondary invagination (Dan and Okazaki, 1956; Gustafson and Kinnander, 1956; Ettensohn, 1985). After basal cell adhesion at the vegetal plate becomes weak, becomes round, and the first invagination occurs (Moore and Burt, 1939; Ettensohn, 1984). The invagination is thought to occur autonomously be caused by four factors; cell growth of the ectodermal layer to cause cell migration at the vegetal side into the blastocoel, growth of cells that compose the archenteron, elongation of the archenteron by rearrangement along the vegetal-animal axis, towing of the gut rudiment by SMCs forming filopodia (Takata and Kominami, 2001; Ettenshon, 1985; Hardin, 1988; Dan and Okazaki, 1956; Gustafson and Kinnander, 1956). However, secondary invagination is no caused by all four factors in all sea urchin species, resulting in species-specific variation, continuous invagination, the cells around the blastopore invaginate continuously without lag phase between the first and second invagination (Ettensohn and Ingersoll, 1992; Kominami and Masui, 1996; Takata and Kominami, 2004). Therefore, sea urchin embryos exhibit species-specific morphologies at the blastula and gastrula stages.

Recently, we studied development of some temnopleurid sea urchins from Japan. We found that the indirect-developing temnopleurid *Temnopleurus toreumaticus* forms a wrinkled blastula with a thick blastocoels wall, whereas other indirect-developing species *T. reevesii, T. hardwickii* and *Mespilia globulus* form smoothed blastulae with a thin blastular wall (Kitazawa et al., 2009, 2010). Embryos of *T. toreumaticus* invaginate continuously to form an archenteron, whereas embryos of *M. globulus* have stepwise invagination (Takata and Kominami, 2004). However, gastrulation of *T. reevesii* and *T. hardwickii* is not fully understood, and in *T. reevesii* development until metamorphosis was described only recently (Kitazawa et al., 2014). Therefore, details of morphogenesis are still unknown.

In this study, we observed blastula and gastrula formation in four temnopleurids using light and scanning electron microscopes to clarify the mechanisms producing the morphological differences among temnopleurids.

## RESULTS

### Internal surface structures of embryos

The internal surface structures of embryos of four temnopleurids were observed by SEM until the mesenchyme blastula stage (Figs 2-5). In *T. toreumaticus*, morulae formed wrinkled blastulae after some cleavages (Fig. 2B-E) and then developed a smoothed surface again (Fig. 2F). During this period, the blastomeres had globular shape and were loosely associated with each other (Fig. 2B,E-). On the surface, there was web of fibers and granular structures of the extracellular matrix (ECM) (Fig. 2B,E’). After disappearance of the wrinkles around 6 h after fertilization, the blastomeres of the blastula started to change shape (Fig. 2F,G). At the vegetal pole, cells around the presumptive PMCs began to extend pseudopod-like structures toward the presumptive PMCs (Fig. 2G) and the ECM expanded on the internal surface except for the area of presumptive PMCs. At this stage, 17.1% of embryos developed this hole-like structure (*n* = 35). After approximately 30 min, this structure became more apparent and it appeared that the ECMs around the presumptive PMCs were covering them by elongation of pseudopod-like structures of the outer cells (Fig. 2H,H-,I). The ratio of embryos with the hole-like structure was 62.5% at this stage (*n* = 48) which then increased to 80.0% (*n* = 15) 7 h after fertilization, and 94.1% (*n* = 17) 7.5 h after fertilization. At 8.5 h after fertilization, the PMCs ingressed into the blastocoel as a mass (Fig. 2J-M).

**Fig. 1.**
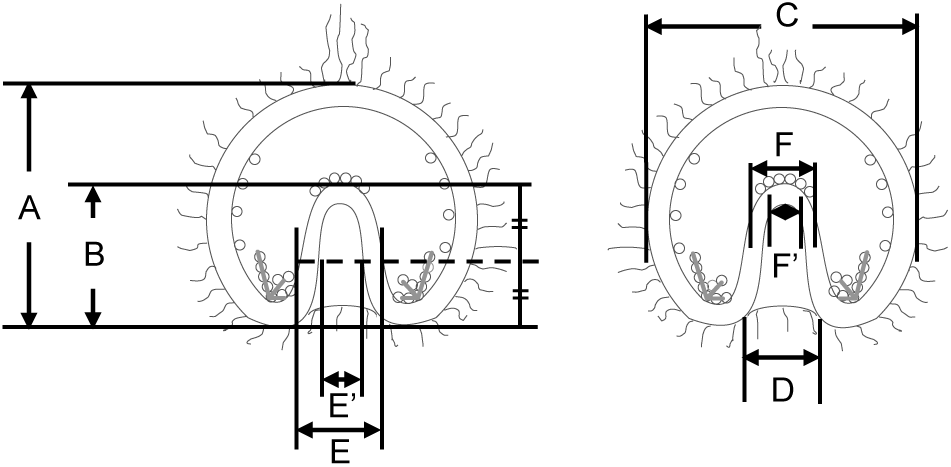
Measurement of gastrulae. Each part of the gastrula was measured; the total length of the embryo (A) or the archenteron (B), the total width of the embryo (C), the diameter of the blastopore (D), the outer or inner diameter of the archenteron at the middle part of the total length of the archenteron (E or E′) or at the tip (F or F′).

**Fig. 2.**
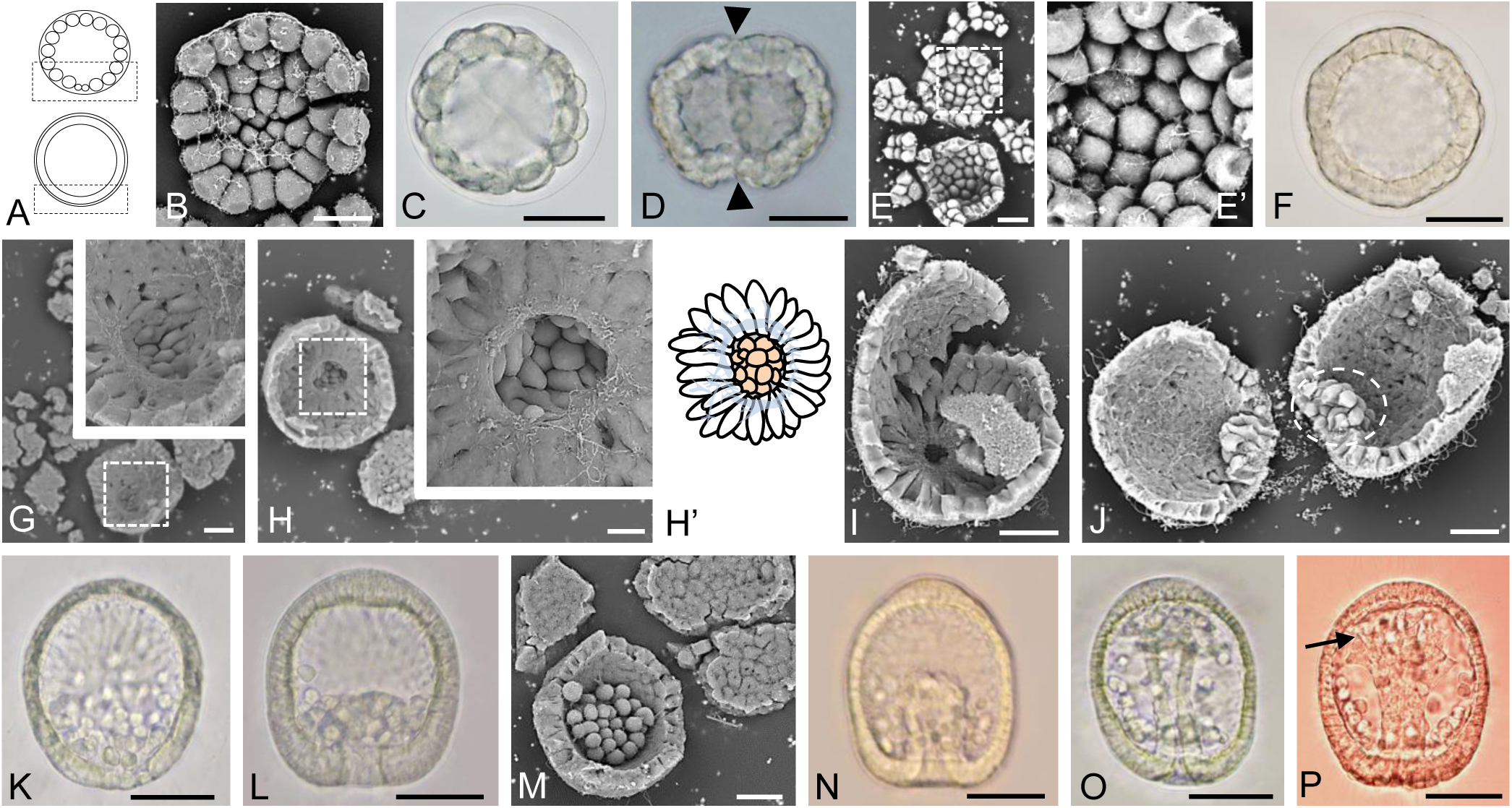
Embryonic development of *T. toreumaticus*. Embryos observed by light (C, D, F, K, L, N–P) or SEM (B, E, E′, G–J, M). **A**. Schematic diagrams of a morula (upper) and blastula (bottom), lateral view. The area inside the black boxes was focused on using SEM. **B**. The internal surface at the vegetal pole of an embryo 3.5 h after fertilization. **C**. A morula 4 h after fertilization. **D, E**. Blastulae 4.5 h after fertilization. The embryos hd developed wrinkles (D; arrowheads). Blastomeres were still orbicular in shape (E, E′) [E′: higher magnification of area inside white box in (E)]. **F, G**. Blastulae 6 h after fertilization. The wrinkles had disappeared (F). At the vegetal pole (G), micromeres-descendants had kept their orbicular shape, but the cells around them had started to produce pseudopod-like structures [insert; higher magnification of the area inside the white box in (G)]. **H**. An embryo 6.5 h after fertilization. As shown in the insert, which show a higher magnification of the area inside the white box, orbicular cells were surrounded by cells that extended pseudopod-like structures toward these cells and there was a ring of ECM at the vegetal pole that seemed to form a hole-like structure. (H′) shows a schematic diagram of the inside of the vegetal plate [center: presumptive PMCs, outer: cells with pseudopod-like structures, blue lines: ECM]. **I**. A hatching blastula 7 h after fertilization. **J**. An embryo 8.5 h after fertilization. PMCs migrated into the blastocoel as a mass (white dashed circle). **K–M**. Embryos at 10 h (K), 11 h (L) or 11.5 h (M) after fertilization [dorso-ventral (K, L, N–P) or animal views (M)]. After migration, PMCs moved into a narrow blastocoel at the vegetal side. **N–P**. Gastrulae 12.5 h (N), 13 h (O) or 15 h (P) after fertilization. The archenteron with the flat tip elongated toward the apical plate and then formed SMCs (N). The middle part of archenteron was narrower (O) and finally the tip attached to the apical plate (P). At this stage, there were some SMCs with filopodia (arrow). Scale bars B, E, G–J, M=20 μm; C, D, F, K, L, N–P=50 μm.

In *T. reevesii* (Fig. 3) and *T. hardwickii* (Fig. 4), each blastomere of morula was adjoined closely (Figs 3B, C, 4B, C). During the blastula stage, the blastomeres extended pseudopod-like structures at the vegetal (Figs 3C, 4B) and lateral sides (Fig. 3E) but the hole-like structure observed in *T. toreumaticus* was not seen (Figs 3G-I, 4C). The blastomeres of *T. hardwickii* developed many filopodia-like structures (Fig. 4B,C). At 7.5-8 h for *T. reevesii* and 9.5-10.5 h after fertilization for *T. hardwickii*, the embryos ingressed PMCs independent to each other (Figs 3I, H, 4D). At this stage in *T. reevesii*, the cells around the PMCs extended pseudopod-like structures toward the PMCs (Fig. 3I).

**Fig. 3.**
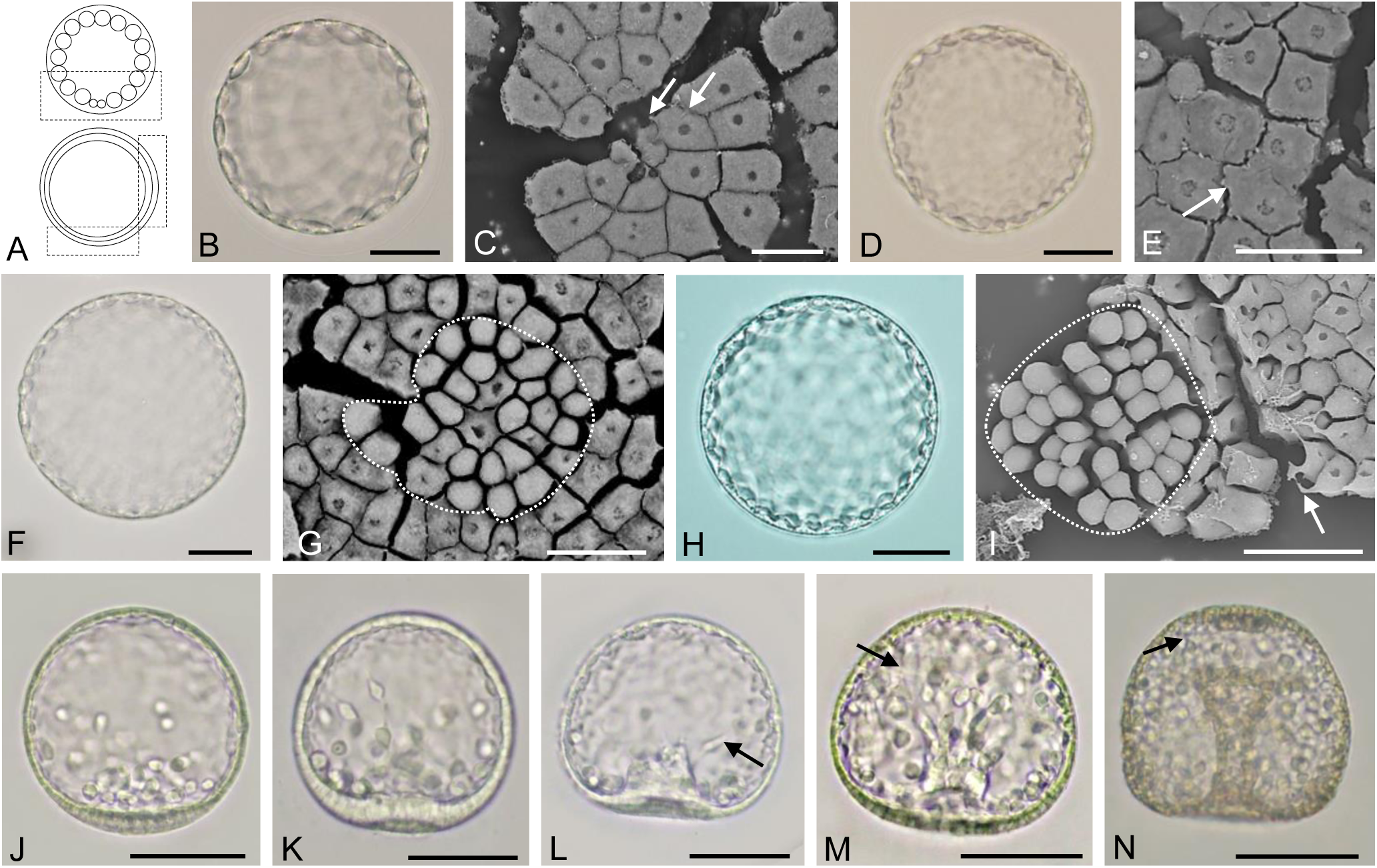
Embryonic development of *T. reevesii*. Embryos observed by light (B, D, F, H, J–M) or SEM (C, E, G, I). **A**. Schematic diagrams of a morula (upper) and blastula (bottom), lateral view. The area inside the black boxes was studied by SEM. **B, C**. Morulae 3.5 h after fertilization. On the internal surface at the vegetal pole, blastomeres were adjoined closely to each other by pseudopod-like structures (white arrows). **D**. An early blastula 4.5 h after fertilization. **E**. The internal surface of the lateral region of an embryo 5.5 h after fertilization. Some cells extended pseudopod-like structures to the neighboring cells. **F, G**. Blastula 6 h after fertilization. By observation of the internal surface of the vegetal plate (G), some presumptive PMCs were globular (area encircled by the dashed white line). **H, I**. Hatching blastulae 7 h after fertilization. By observation of the internal surface of the vegetal plate (I), cells around the globular cells (area encircled by the dashed white line) extended pseudopod-like structures toward this area. **J**. A mesenchyme blastula 11 h after fertilization (lateral view). PMCs were identified at the vegetal area. **K–N**. Gastrulae at 12 h (K), 14 h (L), 15 h (M) or 23 h (N) h after fertilization viewed from the dorso-ventral side. The vegetal plate became thick and then invaginated into the blastocoel (K). SMCs were identified near the tip of the archenteron and some SMCs had filopodia (black arrows) (L, M). The tip of archenteron has a diameter larger than that of the blastopore (N). Scale bars B, D, F, H, J–N=50 μm; C, E, G, I=20 μm.

**Fig. 4.**
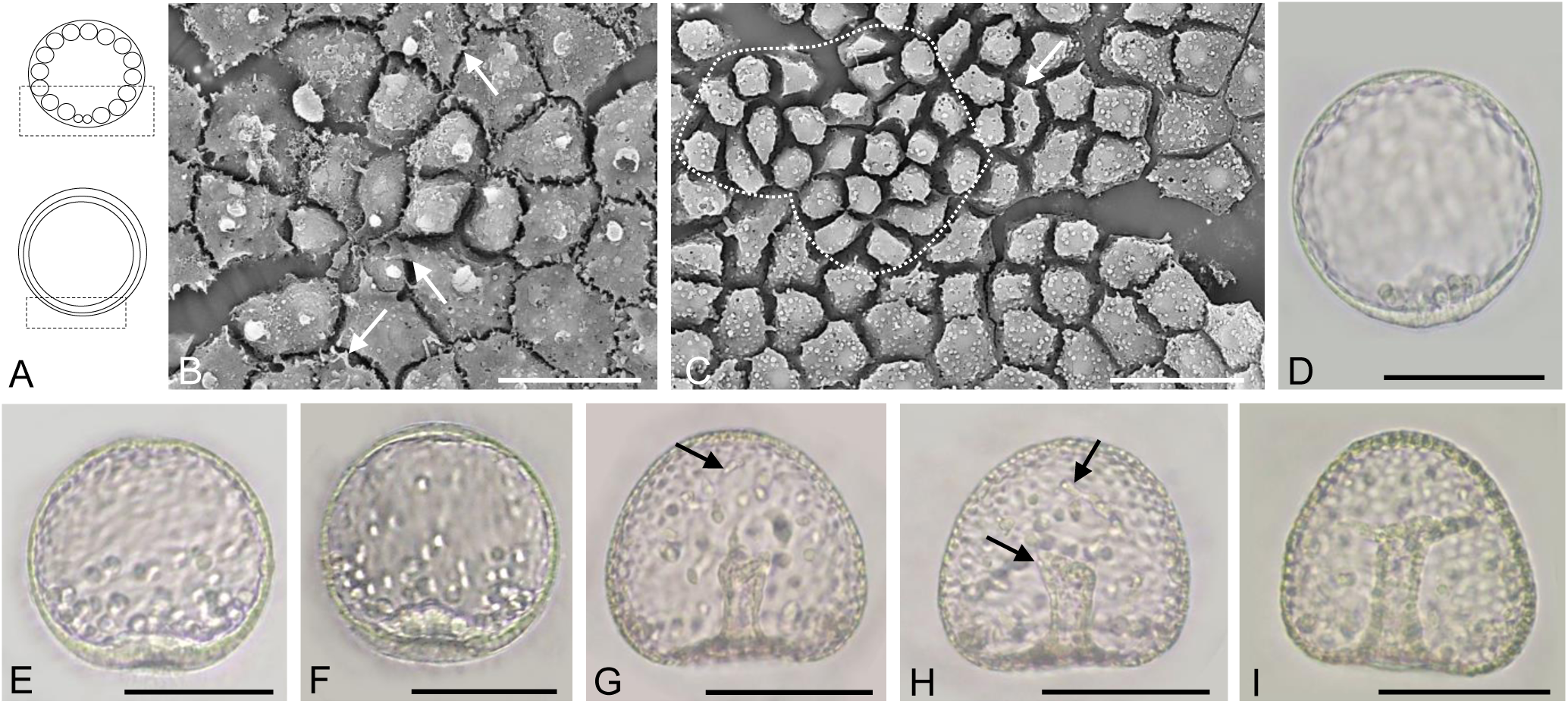
Embryonic development of *T. hardwickii*. Embryos observed by light (D–I) or SEM (B, C). **A**. Schematic diagrams of a morula (upper) and blastula (bottom), lateral view. The area inside the black boxes was examined by SEM. **B**. The internal surface at the vegetal pole of an embryo 4.5 h after fertilization. Each cell extended pseudopod-like structures (white arrows) toward the neighboring cells. **C**. The internal surface at the vegetal pole of an embryo 7 h after fertilization. Some cells had pseudopod-like structures extended toward the adjacent cells (white arrows). Cells enclosed by the dashed white line were identified as the globular cells. **D**. A mesenchyme blastula 11 h after fertilization. **E–I**. Gastrulae at 13 h (E), 14 h (F), 16 h (G), 17 h (H) or 21 h (I) after fertilization (dorso-ventral views). The invagination is slightly curved (E, F) and then the archenteron became thin (G). The length and shape of the archenteron did not change for a while. SMCs occurred and they moved into the blastocoel and formed filopodia (G, H). Finally, invagination was completed (I). Scale bars B, C=20 μm; D–G=50 μm.

**Fig. 5.**
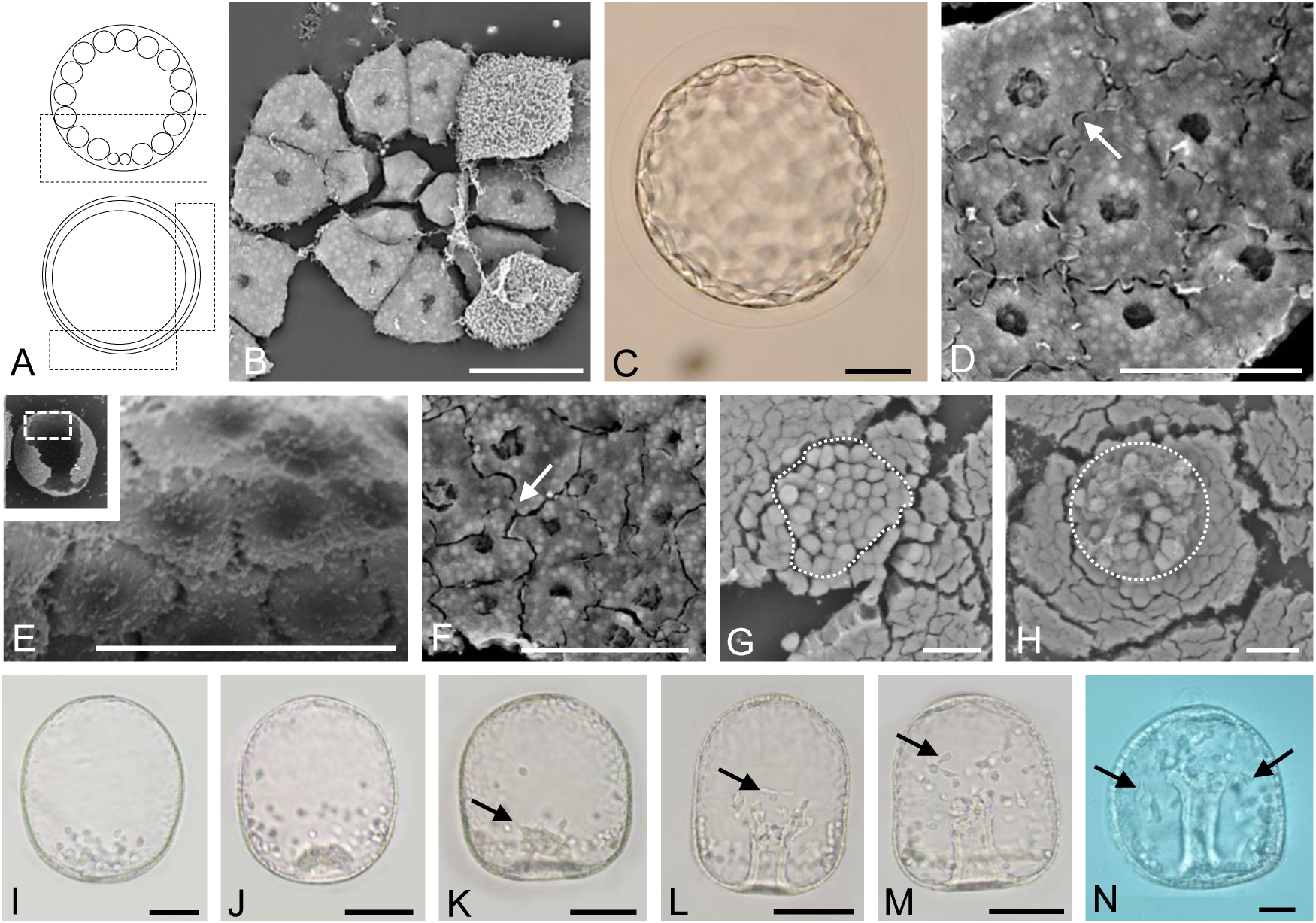
Embryonic development of *M. globulus*. Embryos observed by light (C, I–N) or SEM (B, D–H). **A**. Schematic diagrams of a morula (upper) and blastula (bottom), lateral view. The area inside the black boxes was examined by SEM. **B**. The internal surface structure at the vegetal pole at 3.5 h after fertilization. Each blastomere adjoined closely. **C–E**. Early blastulae 4.5 h after fertilization. Cells at the lateral region with intricate elongated pseudopod-like structures (D). Some specimens had granular structures on the surface of the blastular wall (E: higher magnification of area inside white box in the insert). **F**. The internal surface at the lateral region of an embryo 6 h after fertilization. **G, H**. The internal surface at the vegetal plate of early mesenchyme blastulae 10 h (G) or 11 h (H) after fertilization. In the area enclosed by the dashed white circle are orbicular cells that have started to ingress into the blastocoel as PMCs (H). **I**. A mesenchyme blastula 12 h after fertilization (lateral view). **J–N**. Gastrulae 14 h (J), 15 h (K), 16 h (L), 17 h (M) or 21 h (N) after fertilization (dorso-ventral views). The invaginating vegetal plate is shaped like a hemisphere (J). When the archenteron had invaginated by approximately one fifth of the total length of the embryo (L), the tip of archenteron was flat and released SMCs. Some SMCs formed filopodia (black arrows) and moved in the blastocoel (K–M). Without attachment of the archenteron to the apical plate, the invagination finished (N). Scale bars B, D–G=20 μm; C, H–M=50 μm.

In *M. globulus* (Fig. 5), each blastomere was adjoined closely at the vegetal (Fig. 5B) and lateral sides (Fig. 5D) during the morula and blastula stages (3.5-6 h after fertilization). There were specimens not only with fewer ECMs (Fig. 5D) but also with a lot of ECM granular structures on the internal surface (Fig. 5E). At the lateral side, the blastomeres had different shape compared with those from the vegetal side and the other three species and had many intricate pseudopod-like structures (Fig. 5D,F). However, the ECMs did not develop at the internal surface of the vegetal side and there was no hole-like structure (Fig. 5G,F). At 11.5-12 h after fertilization, the PMCs started to ingress into the blastocoel separately (Fig. 5H).

Table 1 shows a summary of features of blastula in four temnopleurids.

**Table 1.**
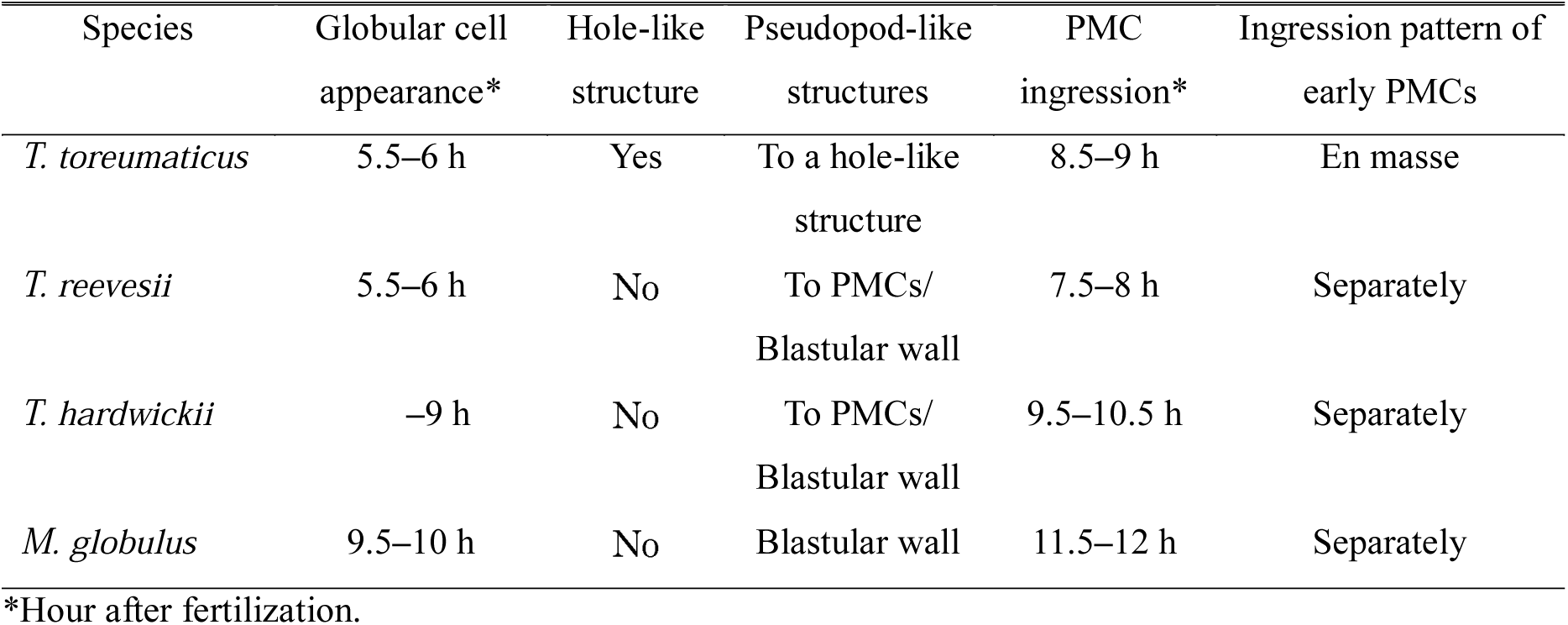
Summary of blastula features of four temnoleurids.

| Species | Globular cell appearance* | Hole-like structure | Pseudopod-like structures | PMC ingression* | Ingression pattern of early PMCs |
| --- | --- | --- | --- | --- | --- |
| *T. toreumaticus* | 5.5–6 h | Yes | To a hole-like structure | 8.5–9 h | En masse |
| *T. reevesii* | 5.5–6 h | No | To PMCs/<br>Blastular wall | 7.5–8 h | Separately |
| *T. hardwickii* | –9 h | No | To PMCs/<br>Blastular wall | 9.5–10.5 h | Separately |
| *M. globulus* | 9.5–10 h | No | Blastular wall | 11.5–12 h | Separately |

2 \*Hour after fertilization.

### Manner of invagination of the archenteron

The results of gastrulation in *T. toreumaticus* basically support the findings of Takata and Kominami (2004). The vegetal plate became thicker with PMCs ingression 10 h after fertilization (Fig. 2K). After 1 h, the vegetal plate ingressed into the blastocoel by 22.6 ± 4.3% of the total length of the embryo (Figs 2L, 6A, B). At 12 h after fertilization, the archenteron ingressed by 44.9 ± 8.6% and the diameter was narrower than in the previous hour (Fig. 2N). At approximately 70% of ingression, the middle part of the archenteron became narrower and the wall of the archenteron became thinner (Fig. 2O). At this stage, SMCs were identified at the tip of the archenteron. At 15 h after fertilization, the tip of the archenteron was attached to the apical plate (Fig. 2P) and the invagination ratio was 77.1 ± 7.4% (as the ectodermal area at the apical plate was included in the total length of the embryo in this experiment, the ratio is not 100%) (Fig. 6B). These results indicate that this species has continuous invagination without a lag phase.

**Fig. 6.**
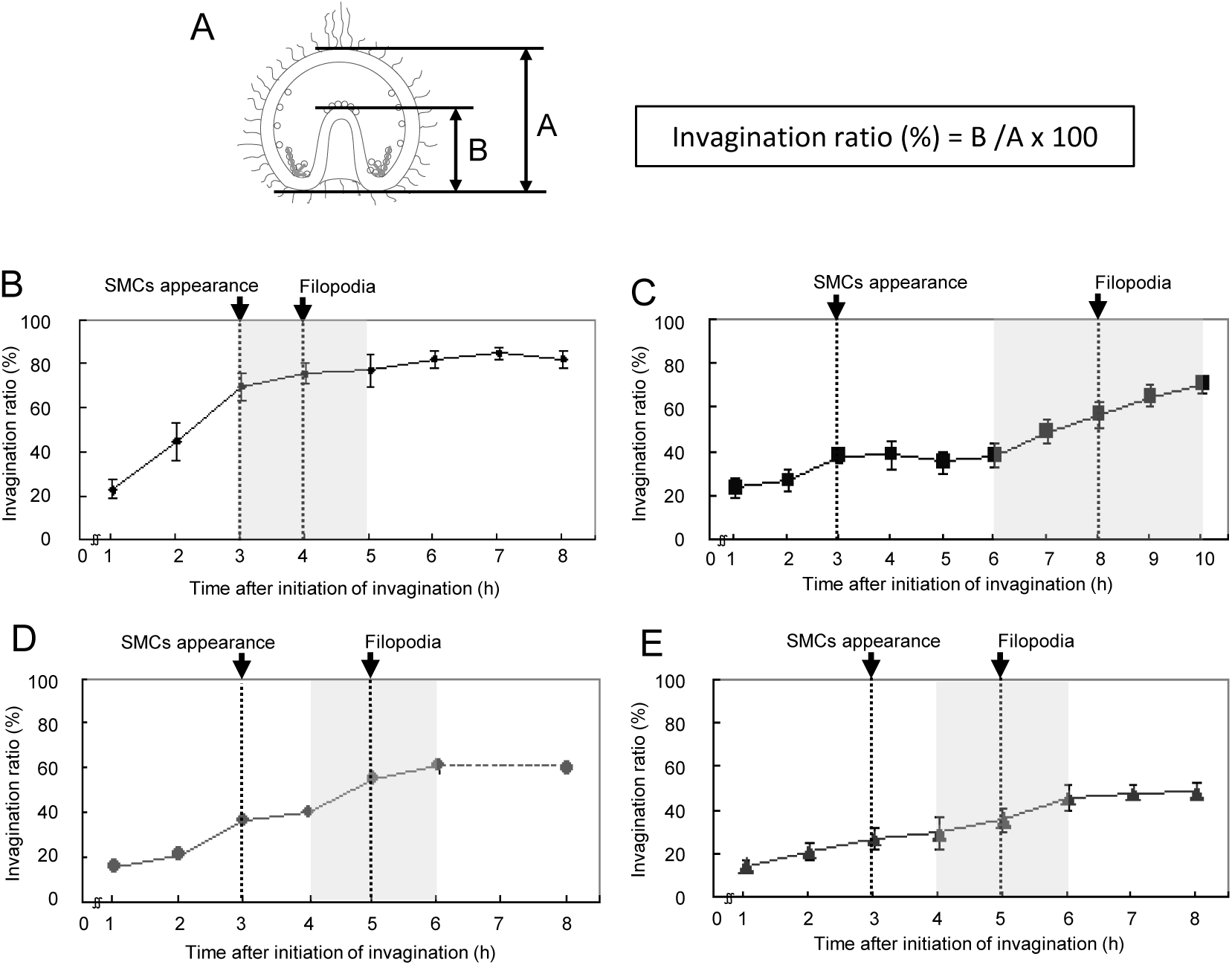
Pattern of invagination of the archenteron in four temnopleurids. **A**. A schematic diagram of a gastrula derived from Fig. 1 for measurements to find the invagination ratio. **B**. *T. toreumaticus.* **C**. *T. reevesii.* **D**. *T. hardwickii*. **E**. *M. globulus.* Y-and X-axes show the invagination ratio (%) or the time after initiation of invagination. Gray areas show the secondary invagination. The timing of SMCs appearance or SMCs with filopodia show the stage when over 60% of specimens have these features. Bars: SDs.

In *T. reevesii*, PMCs ingressed into the blastocoel 11 h after fertilization (Fig. 3J). The vegetal plate became thicker and started to ingress slightly (Fig. 3K). At 14 h after fertilization, the archenteron ingressed until 23.8 ± 4.6% and SMCs occurred at the tip of the archenteron (Figs 3L, 6C). The archenteron ingressed by 38.2 ± 3.4% in 2 hours, but the ratio did not change (Fig. 6C). At 20 h after fertilization, the archenteron ingressed again until approximately 50% (Fig. 6C). The tip of the archenteron did not attach to the apical plate and invagination of the archenteron finished 23 h after fertilization (Fig. 3N). Embryos of *T. hardwickii* had similar invagination pattern of *T. reevesii* (Fig. 4). After PMC ingression 11 h after fertilization (Fig. 4D), the vegetal plate became thicker and started to invaginate slightly 13 h after fertilization (Fig. 4E). The archenteron ingressed keeping a smoothed curve shape and the PMCs started to move into the blastocoel (Fig. 4F). At 14 h after fertilization, the invagination ratio of the archenteron was 15.9 ± 2.6% and then increased to 36.2 ± 2.7% in 2 hours (Figs 4G, 6D). Around this stage, some SMCs started to move from the tip of the archenteron into the blastocoel (Fig. 4G, H). At 17 h after fertilization, the invagination ratio was still 39.8 ± 2.4% (Fig. 6D). Finally, the invagination finished at approximately 60% (19 h after fertilization) without attachment of the archenteron to the apical plate (Figs 4I, 6D).

In *M. globulus*, the PMC ingression started 11 h after fertilization (Fig. 5G-I) and then 3 h later invagination of the archenteron began (14.4 ± 2.8%) (Figs 5J, 6E). In most developing specimens, the archenteron ingressed by 20.6 ± 3.9% 15 h after fertilization (Fig. 6E) with SMCs at the tip (Fig. 5K). One hour later, the invagination ratio increased to 27.0 ± 5.2% (Fig. 6E) and some SMCs formed filopodia and started to move into the blastocoel (Fig. 5L). After passing the lag phase 16-17 h after fertilization (Figs 5M, 6E), the archenteron elongated suddenly and then finished invagination at 19 h after fertilization (48.4 ± 4.1%) (Figs 5N, 6E).

### Morphological changes of the archenteron

In our study of temnopleurid gastrulation, each region of an embryo was measured (Fig. 1). At first, the diameter of the blastopore for the total width of the embryo was measured (Fig. 7). In *T. toreumaticus*, the ratio was 33.5 ± 9.0% at initiation of invagination until just before the finish of invagination and then decreased to 9.3 ± 6.5% 18 h after fertilization (Fig. 7B). The diameter did not change in *T. reevesii* during invagination (approximately 30%, Fig. 7C). In *T. hardwickii*, the diameter of the blastopore was constant until the end of first invagination (31.5 ± 5.7%) but then decreased after the second invagination started (25.7 ± 4.3%) (Fig. 7D). In *M. globulus*, the diameter was constant until the end of the lag phase (29.4 ± 4.0% 17 h after fertilization). However, it decreased after the initiation of the secondary invagination (23.4 ± 4.3%) (Fig. 7E).

**Fig. 7.**
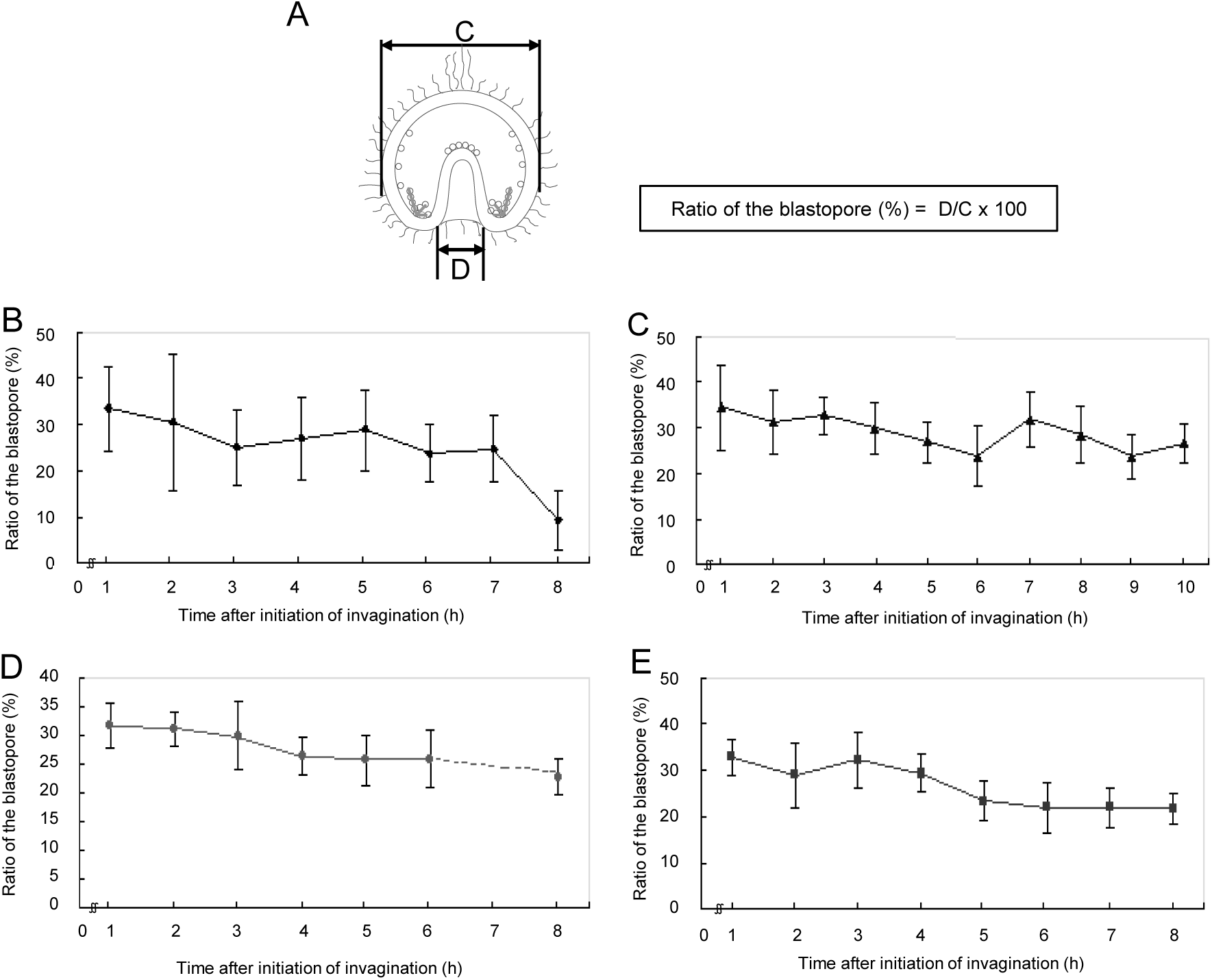
Ratio of the diameter of blastopore to the total width of embryo in four temnopleurids. **A**. A schematic diagram of a gastrula for measurement to find the ratio of the blastopore. **B**. *T. toreumaticus.* **C**. *T. reevesii.* **D**. *T. hardwickii*. **E**. *M. globulus.* Y-and X-axes show the ratio of the blastopore (%) or the time after initiation of invagination. Bars: SDs.

Next, the archenteron diameter was measured at the middle part (Figs 1, 8A). In *T. toreumaticus*, the archenteron diameter had decreased 2-3 h after initiation of invagination (28.5 ± 5.0 to 18.9 ± 2.4 μm) and did not change thereafter (Fig. 8B). To determine whether this decrease was caused by a decrease in thickness of the cells at the archenteron wall, the thickness of the archenteron wall was calculated (Fig. 8A). During invagination, the thickness of the archenteron wall was 8.7 ± 2.7 μm until 2 h after initiation of invagination and then decreased to 5.8 ± 1.8 μm 1 h later (Fig. 8C). Therefore, the decrease in thickness of the archenteron wall at both sides was approximately 6 μm and this decrease may have caused a decrease in the diameter of the archenteron. In *T. reevesii*, the diameter of the archenteron decreased as secondary invagination progressed (from 25.2 ± 4.5 μm 14-18 h after fertilization to 16.9 ± 2.7 μm 23 h after fertilization) (Fig. 8D). The thickness of the archenteron wall was 5.3 ± 2.4-6.0 ± 1.9 μm until just after the initiation of the secondary invagination but it decreased as secondary invagination progressed (Fig. 8E). This means that the decrease in diameter of the archenteron is caused solely by decrease in thickness of the archenteron wall. In *T. hardwickii*, it was difficult to identify each area before the end of the first invagination (Fig. 4G). At this stage, the diameter of the archenteron was 19.7 ± 1.7 μm and then slightly decreased to 17.1 ± 1.8 μm just after the initiation of the secondary invagination (Fig. 8F). In this species, the thickness of the archenteron wall did not change during invagination (Fig. 8G). In *M. globulus*, it was difficult to identify each area before the lag phase and the diameter of the archenteron was 31.7 ± 4.5 μm (Fig. 8H). From the initiation of the secondary invagination, it decreased to a constant diameter of 23.6 ± 3.3 μ m. The thickness of the archenteron wall was 8.5 ± 1.8 μm at 17 h after fertilization, and then decreased to 6.3 ± 1.4 μm 1 hour later (Fig. 8I). This means that the decrease in the diameter of the archenteron is caused not by a decrease in the thickness of the archenteron wall only.

**Fig. 8.**
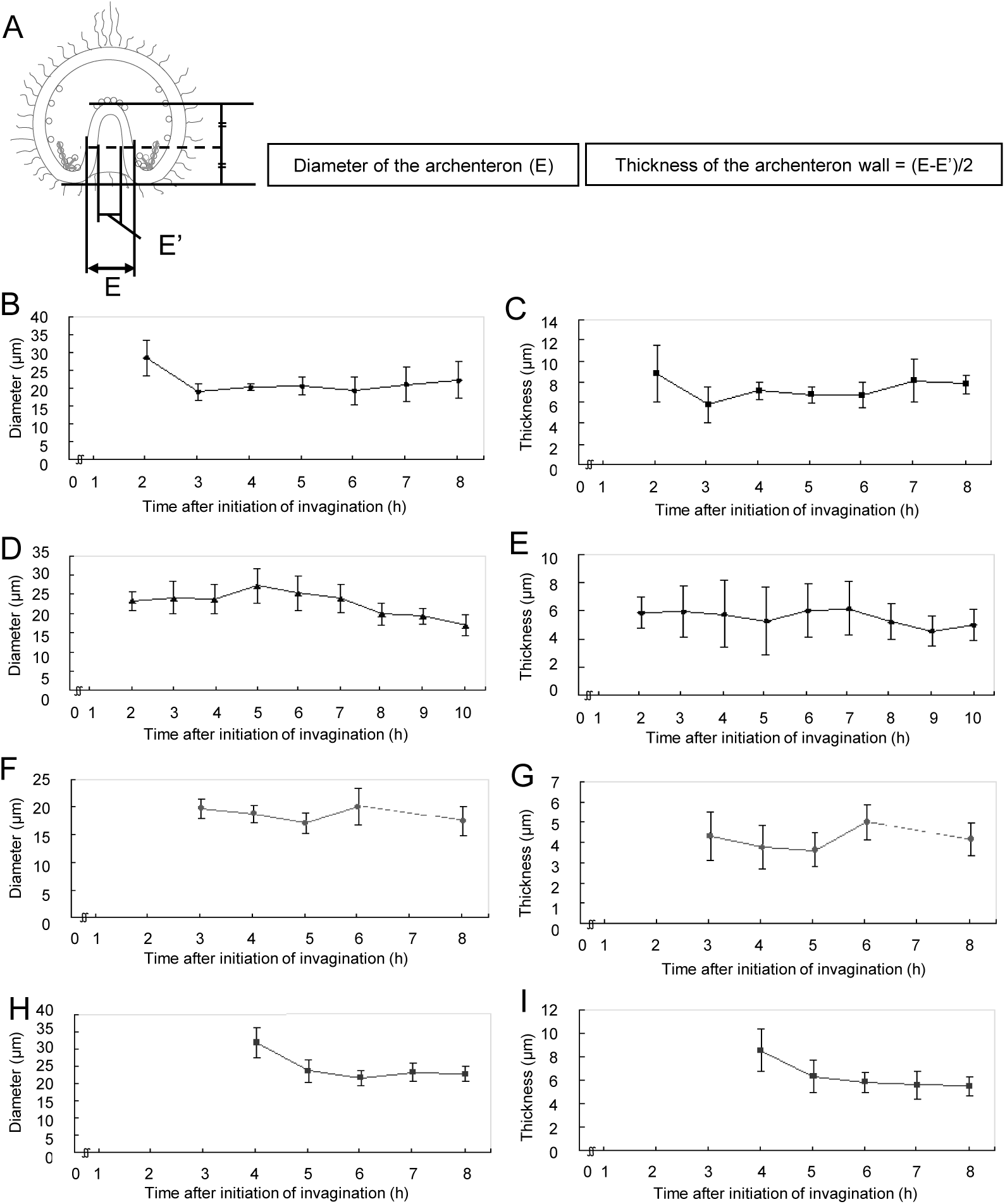
Diameter of the archenteron and thickness of the archenteron wall in four temnopleurids. **A**. A schematic diagram of a gastrula for measurement of the diameter of the archenteron and measurement to find the thickness of the archenteron wall. **B, C**. *T. toreumaticus.* **D, E**. *T. reevesii.* **F, G**. *T. hardwickii*. **H, I**. *M. globulus.* Y-and X-axes show the diameter of the archenteron (B, D, F, H) and thickness of the archenteron wall (C, E, G, I) or the time after initiation of invagination. Bars: SDs.

In *T. toreumaticus*, the outer and inner diameters of the archenteron on the tip were constant during invagination of the archenteron (Fig. 9B,C). In *T. reevesii*, the outer and inner diameters were 27.5 ± 4.8 and 19.1 ± 6.2 μm, respectively, just after the start of the first invagination, but decreased 1 h later (Fig. 9D,E). The diameters stayed constant until secondary invagination and then increased to 33.5 ± 5.7 and 21.3 ± 4.4 μm 23 h after fertilization. In *T. hardwickii*, the outer and inner diameters were 23.7 ± 3.3 and 12.7 ± 3.3 μm respectively at the start of secondary invagination and then increased to 28.7 ± 3.6 and 14.5 ± 3.8 μm (19 h after fertilization) (Fig. 9F,G). However, the diameters decreased. In *M. globulus*, the outer and inner diameters decreased as invagination progressed [33.9 ± 5.9 and 21.3 ± 4.8 μm, respectively, at the initiation of invagination (Fig. 9H); 27.3 ± 3.7 and 11.8 ± 3.3 μm at the end of the invagination (Fig. 9I)].

**Fig. 9.**
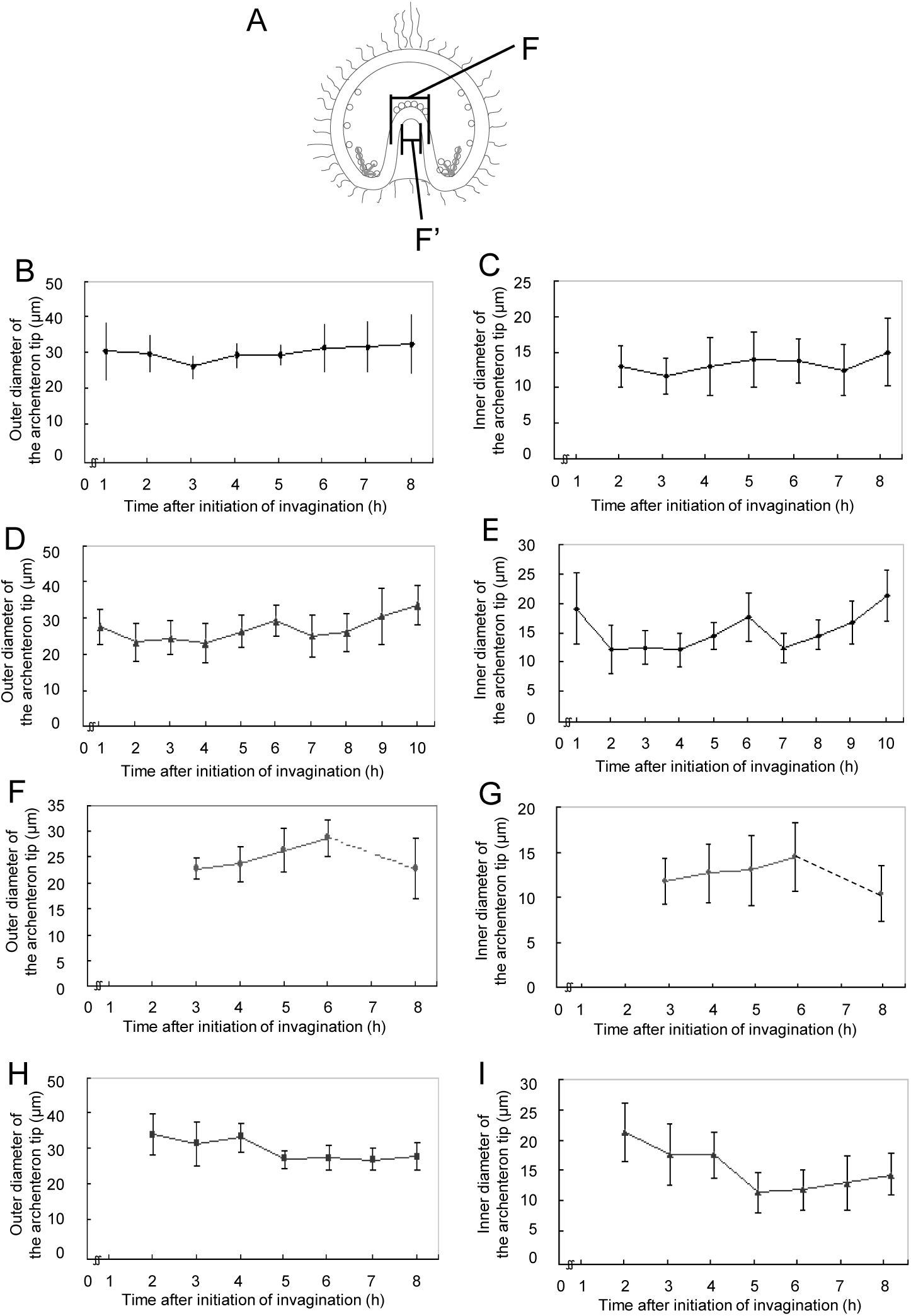
The diameter of the tip of the archenteron in four temnopleurids. **A**. A schematic diagram of a gastrula for measurement of the outer and inner diameter of the tip of the archenteron. **B, C**. *T. toreumaticus.* **D, E**. *T. reevesii.* **F, G**. *T. hardwickii*. **H, I**. *M. globulus.* Y-and X-axes show the outer (B, D, F, H) and inner diameter of the tip of the archenteron (C, E, G, I) or the time after initiation of invagination. Bars: SDs.

The ratio of the internal archenteron length without the wall of the tip for the total length of the embryo was as the invagination ratio of the internal archenteron. In *T. toreumaticus, T. hardwickii* and *M. globulus*, this ratio became higher as invagination progressed, while in *T. reevesii*, this ratio was very similar to the ratio of invagination of the archenteron at 2 h after initiation of invagination (25.4 ± 3.6% and 26.7 ± 5.1%) (31.3 ± 8.8% and 44.9 ± 8.6% 12 h after fertilization in *T. toreumaticus;* 11.1 ± 3.9% and 21.3 ± 2.4% 16 h after fertilization in *T. hardwickii;* 12.0 ± 4.8% and 20.6 ± 3.9% 16 h after fertilization in *M. globulus).* The difference between the invagination ratio of the internal archenteron length and the archenteron length off the whole embryonic length was always about 8.0 ± 2.0% 3 h after the initiation of invagination.

**Timing of appearance and filopodia formation of SMCs**

The SMCs in four temnopleurids appeared approximately 3 h after initiation of invagination (Fig. 6; 70.0% of *T. toreumaticus*, 65.0% of *T. reevesii*, 80.0% of *T. hardwickii* and *M. globulus).* After this stage, a part of the SMCs formed filopodia and the number of specimens with SMCs with one or more filopodia was counted. In *T. toreumaticus*, 65.0% of specimens had SMCs with filopodia before 1 h at the end of invagination (Fig. 6). Embryos of the three other species *(T. reevesii, T. hardwickii* and *M. globulus)* with stepwise invagination had SMCs with filopodia during the secondary invagination (Figs 2-6) [85.0% in *T. reevesii* 8 h after initiation of invagination (21 h after fertilization); 75.0% of *T. hardwickii* 4 h after initiation of invagination (18 h after fertilization); 70.0% in *M. globulus* 5 h after initiation of invagination (18 h after fertilization)].

## DISCUSSION

In this study, structures of the blastulae and gastrulae indicated species-specific features among four temnopleurids (Figs 2-5). The shape and position (vegetal or lateral) of blastomeres in blastulae differed among species. The loosely adjacent blastomeres on the blastular wall of *T. toreumaticus* had a globular shape until the wrinkled blastula stage (Fig. 2). In *T. toreumaticus* the cell number at hatching is only 500, whereas in other species it is 600-800 (Masuda, 1979). As the diameter of the blastulae is similar among these species (Kitazawa et al., 2010), we suggest that fewer cleavages cause the difference in the cell shape and adhesion among blastomeres at the same developmental stage.

The blastulae of the temnopleurids studied have blastomeres with different kinds of pseudopod-like structures (Figs 2-5). In *T. toreumaticus*, the blastomeres around the presumptive PMCs extended pseudopods to them (Fig. 2G,H). Immers (1961) indicated that in sea urchins the formation of filopodial projections of mesenchyme cells is supported by a matrix of sulfated polysaccharides combined with proteins. The differences in the shape of the pseudopod-like structures among temnopleurids may be caused by different substances within the matrix. Recently, Yaguchi et al. (2015) indicated that adhesion between blastomeres at the early cleavages was very loose, and that the blastomeres had many protrusions attached to the outer ECM and the hyaline layer at the cleavage furrow in *T. reevesii.* Also, it was difficult to divide each blastomere of the early cleavage-staged embryos of *T. hardwickii* and *M. globulus* because of the outer ECM and the hyaline layer (data not shown). However, in our observations of blastulae, the space between blastomeres became narrow and connected by complex pseudopod-like structures (Fig. 3). The blastomeres around the presumptive PMCs seemed to form layers by elongated pseudopod-like structures (Fig. 3I). These phenomena indicate that adhesion between blastomeres changes from loose to tight. Future work should analyze whether the early protrusions form the pseudopod-like structures at the blastula stage.

The distribution of the ECMs was different among species and position (vegetal or lateral). The blastocoelic surface develops many kind of ECMs that include mucopolysaccharides (Okazaki and Niijima, 1964), glycoproteins fibronectin, and laminin (Spiegel et al., 1983; Benson et al., 1999), and collagen (Kefalides et al., 1979; Crise-Benson and Benson, 1979). Furthermore, in the basal lamina distribution of fibronectin and laminin differ among species (Spiegel et al., 1983; Katow et al., 1982). The blastocoelic surface of the vegetal plate of *T. toreumaticus* had was a hole-like structure in the ECM (Fig. 2G,H). This was absent in the other species (Figs 3-5). In *Lytechinus variegatus*, the blastocoelic surfaces were covered with a thin basal lamina composed of fibrous-and non-fibrous materials before PMC ingression and then a web-like ECM became located at the animal hemisphere (Galileo and Morrill, 1985). Galileo and Morrill (1985) found that the blastoemeres, before hatching on the blastocoel wall, were intertwined and had patchy meshwork of ECM. They were connected by thin cellular processes to each other perpendicular to the animal-vegetal axis. In addition, the blastocoel wall around the animal hemisphere developed a hole without ECM. Recently it was reported that the presumptive PMCs lose laminin distribution related with the gene regulatory network sub-circuit for basal lamina remodeling that include *tbr*, *dri* and *hex* by knockdown of these genes (Saunders and McClay, 2014). Therefore, we suggest that the hole-like structure observed in *T. toreumaticus* may be formed of laminin added by these genes, and that embryos of the temnopleurids in our study may have different amount and distribution of laminin. Examination of the photographs published by Amemiya (1989) of *Hemicentrotus pulcherrimus* and *Pseudocentrotus depressus* revealed that the blastomeres of the animal side elongate to the vegetal pole. However, the embryos did not form a vegetal hole in the ECM. Furthermore, Amemiya (1989) reported that PMC patterning is caused by accumulation of fibrils of the basal lamina. This report supports our findings that the ECM hole of *T. toreumaticus* may attach PMCs at the vegetal plate. The fixative used in our study was very similar to method lacking calcium ions in Amemiya (1989). Therefore, our results may indicate that there are differences in the distribution of a calcium-dependent ECM among species and that *T. toreumaticus* has a lot of calcium-dependent ECM or calcium-independent ECM.

In *M. globulus*, each blastomere of the blastula became located close to the surface along with filament structures around the 8th cleavage. A sheet-like structure of mucopolysaccharides on the blastocoelic surface was identified for movement of SMCs on these structures (Endo and Uno, 1960). In our study, we did not identify well developed filamentous structures in blastulae of *M. globulus* but observed some ECMs on the blastocoelic surface (Fig. 5E). Endo (1966) reported that in *Mespilia* when the PMCs ingress, they dispose of their apical cytoplasm while still attached by desmosomes to neighboring cells. On the other hand, in *Arbacia* (Gibbins et al., 1969) and *Lytechinus pictus* (Katow and Solursh, 1980), the desmosomes disappear from the PMCs. Therefore, analysis of the ultrastructure of the blastular wall among these temnopleurids is needed in the future.

The ECMs is important for cell movement, such as PMC migration for PMC differentiation, modulation of epithelial cell polarity, and gastrulation (Solursh and Lane, 1988; Katow et al., 1982; Fink and McClay, 1985; Amemiya, 1989; Adelson and Humphreys, 1988; Ingersoll and Ettensohn, 1994; Berg et al., 1996). In *L. pictus*, PMCs have six types of cell processes, depending on a specific component of the basal lamina substratum, that are involved in cell migratory behavior (Katow and Solursh, 1981). In our study, the PMCs of *T. reevesii*, *T. hardwickii* and *M. globulus* ingressed separately. We suggest that in *T. toreumaticus*, PMC ingression en masse may be caused by inner ECM distribution at the vegetal plate. We also observed some kind of process structures (Figs 2-5) and suggest that these structures may cause cell movement.

Gastrulation in *T. reevesii*, and *T. hardwickii* and *M. globulus* is by stepwise invagination with a lag period (Figs 3-6). This means that that the mechanisms of gastrulation in these temnopleurids may be different to that of *T. toreumaticus* with continuous invagination. Although timing of the initiation of invagination in *T. toreumaticus* and *M. globulus* in our study was different from that reported by Takata and Kominami (2004), it suggests that differences between batches of embryos or geographic location [Yamaguchi area in the present study; Kouchi and Ehime areas in Takata and Kominami (2004)] of the same species may cause different developmental speeds of embryos.

The mechanisms of invagination of the archenteron of each species were considered according to four factors: pushing of the vegetal cells into the blastocoel by cell growth at the animal pole (Takata and Kominami, 2001), elongation of the cells forming the archenteron, re-arrangement of the cells forming the archenteron along the animal-vegetal axis and toughening of the gut rudiment by the filopodia of the SMCs ingressed into the blastocoel from the tip of the archenteron (Ettenshon, 1985; Hardin, 1988; Dan and Okazaki, 1956; Gustafson and Kinnander, 1956). In *T. toreumaticus*, the blastopore became narrow at the end of invagination (Fig. 7B). The blastopore of *Scaphechinus milabilis*, which has continuous invagination becomes narrow around the end of invagination (Kominami and Masui, 1996). Therefore, we suggest that the archenteron of *T. toreumaticus* may also elongate by continuous ingression of the cells around the blastopore into the blastocoel. In irregular sea urchins with continuous invagination, the diameter of the archenteron during invagination does not change so that the cell rearrangement does not affect elongation of the archenteron (Kominami and Masui, 1996; Takata and Kominami, 2004). However, we observed that in *T. toreumaticus* the diameter at the mid archenteron and the thickness of the archenteron wall decreased rapidly. There is a possibility that the rearrangement and elongation of the archenteron cells causes elongation of the archenteron. In *T. toreumaticus* and irregular sea urchins, the early developmental events including PMCs ingression, initiation of the invagination of the archenteron, and formation of the SMCs, start and finish at relatively early stages. This suggests that developmental events may be accelerated and omitted overall, and then involved the continuous invagination of the archenteron. In this species the SMCs and elongated filopodia formed near the end of invagination, which means that the SMCs may not toughen the archenteron. The final degree of invagination is about 94% (Fig. 6B) and the primary pore canals of this species do not maintain the body width (Kitazawa et al., 2014). We suggest that the continuous invagination occurs by elongation of the archenteron itself.

In *T. reevesii*, the thickness of the archenteron wall decreased after the lag period (Fig. 6C). This suggests that cell elongation causes elongation of the archenteron. The diameter and thickness of the archenteron wall at the mid archenteron decreased. This suggests that another decrease was caused by rearrangement of the cells in the archenteron (Fig. 8D,E). It is a possible that the SMCs cause elongation of the archenteron because of the long lag period and period for secondary invagination in this species (Fig. 6C), and SMCs formed filopodia during secondary invagination. After initiation of the invagination, the thickness of the archenteron tip became thin temporally in only this species (data not shown).

In *T. hardwickii*, the diameter of the blastopore decreased from the end of the first invagination to the initiation of the secondary invagination (Fig. 6D). As *S. milabilis* (Kominami and Masui, 1996), results suggest that this decrease of *T. hardwickii* may be caused by growth at the animal pole pushing the cells at the vegetal pole causing elongation the archenteron. The diameter at the mid archenteron decreased during the lag period (Fig. 8F), caused by rearrangement of the cells of the archenteron along the animal-vegetal axis. The thickness of the archenteron wall was constant (Fig. 8G) and it may mean that elongation of the archenteron is caused not by cell elongation, but by rearrangement of the cells. SMCs with filopodia were observed during secondary invagination, and it is possible that the SMCs cause elongation of the archenteron. However, invagination in *T. hardwickii* finished at about 60% of the whole embryonic length and SMCs were only identified near the end of the invagination. Therefore, it is possible that the SMCs with filopodia do not cause elongation of the archenteron but toughen the tip of the archenteron at the presumptive oral region.

In *M. globulus*, the cells of the archenteron became thin from the end of the lag period to the initiation of the secondary invagination (Fig. 8H,I). Takata and Kominami (2004) reported that in *M. globulus* the rearrangement was not remarkable. The tip of the archenteron did not attach to the apical plate, nor did SMCs disperse into the blastocoel. Furthermore, the cell number of the archenteron did not change. Therefore, it is suggested that elongation of the cells to form the archenteron causes elongation of the archenteron. The diameter of the blastopore decreased during invagination (Fig. 7E) which means that the push of the vegetal cells by growth of the animal cells may cause elongation of the archenteron in this species.

Formation of SMCs occurred at the same time in the four temnopleurids studied. However, the three species with stepwise invagination needed a longer invagination period than the species with continuous invagination and their final invagination ratio was approximately 60% (Fig. 6). Their SMCs formed filopodia during the late invagination period (Fig. 6) and it is possible that the SMCs change the direction of elongation of the archenteron to the presumptive oral region by toughing the tip of the archenteron. In addition, Amemiya et al. (1982) reported that the pseudopodia from the SMCs may pull up the archenteron in *H. pulcherrimus* and *P. depressus* but not in *A. crassispina* because it does not form many pseudopodia.

The diameters at the tip of the archenteron indicate different changes among species (Fig. 9). These results indicate that the feature at the tip of the archenteron may cause species-specific invagination or the formational process of the coelomic pouches. This conclusion is supported by findings that the pattern of formation of the primary pore canal from the coelomic pouches is different among these species (Kitazawa *et al.*, 2012, 2014).

Our results indicate that temnopleurids have species-specific differences during early morphogenesis, including blastula formation and invagination of the archenteron including effective factors in the same family (Table 1, Fig. 10). *Temnopleurus toreumaticus* develops some species-specific features like wrinkled egg and wrinkled blastula formation at the early developmental stages (Kitazawa et al., 2009, 2010). Phylogenetic analysis based on allozyme data of temnopleurids suggest that *T. toreumaticus* and *T. reevesii* are more closely related than *T. hardwickii* and *M. globulus* (Matsuoka and Inamori, 1996). However, based on morphological and molecular analysis, Jefferies et al. (2003) determined that *M. globulus* and *T. reevesii* are more closely related to each other than to *T. toreumaticus*. Recently, we also observed development of another temnopleurid, *Temnotrema sculptum* and this species is very similar to *T. reevesii*, *T. hardwickii* and *M. globulus*, but not *T. toreumaticus* (Fujii et al., 2015). Therefore, we hypothesize that after divergence, *T. toreumaticus* evolved more species-specificities at early developmental stages than other temnopleurids.

**Fig. 10.**
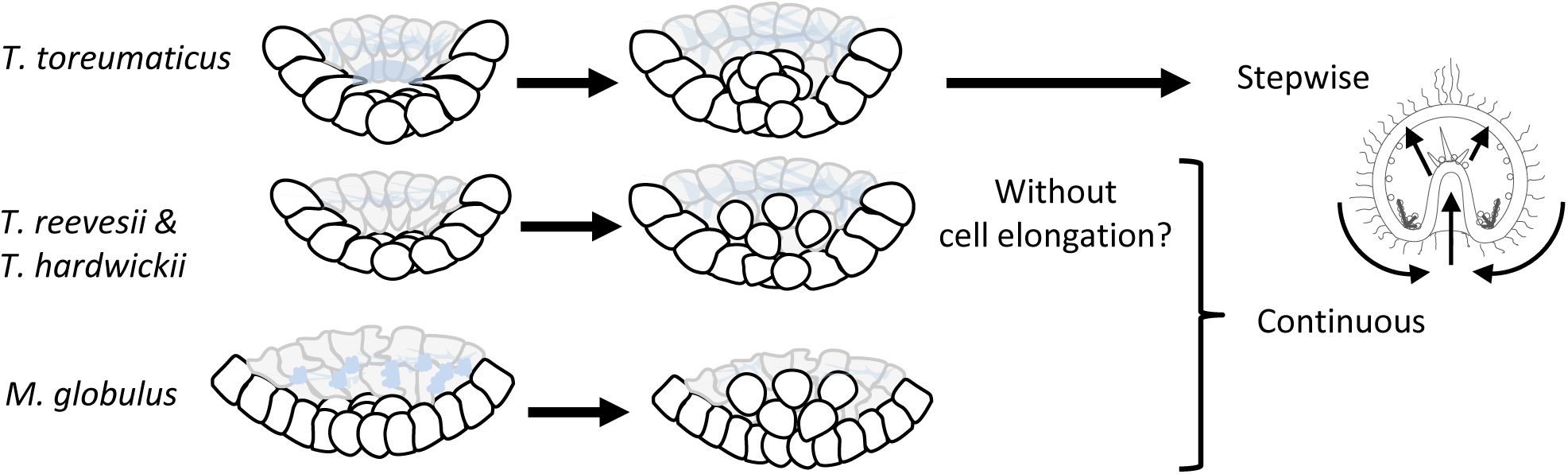
Summary of blastula and gastrula formation in four temnopleurids.

## MATERIALS AND METHODS

### Spawning and embryonic culture

Adult *Temnopleurus toreumaticus*, *T. reevesii*, *T. hardwickii* and *Mespilia globulus* were collected from the Inland Sea (Setonai), Yamaguchi Prefecture, Japan. They were induced to spawn by injection of a small amount of 0.5 M KCl solution into the body cavities from June to December for *T. toreumaticus*, June to October for *T. reevesii*, April to November for *T. hardwickii*, and August to December for *M. globulus*. The eggs were washed with filtered sea water (FSW) and then fertilized. The fertilized eggs were transferred into a glass dish filled with artificial sea water (ASW; TetraMarin^®^ Salt Pro, Tetra, Melle, Germany) and cultured at 24°C. The embryos were observed under a microscope (OPTIPHOT-2, Nikon, Tokyo, Japan) and photographed using digital cameras (FinePix F710, Fujifilm, Tokyo, Japan; F200EXR, Fujifilm; µ810, Olympus, Tokyo, Japan).

### Fixation and observation of embryos

For SEM observation, embryos were fixed, dehydrated and mounted on aluminum stubs using double-sided conductive aluminum tape according to Kitazawa et al. (2012). After dividing the embryos with a hand-held glass needle, the specimens were coated with gold using a fine ion sputter coater (E-1010, Hitachi High-Technologies, Tokyo, Japan), observed and photographed under a scanning electron microscope (Miniscope TM-1000S, Hitachi).

For observation of gastrulation, after 10 h after fertilization embryos were fixed in 4% formalin ASW for approximately 45 min every hour. Fixed embryos were exchanged gradually from 70% ethanol to ion-exchanged water in a 96-well plastic plate coated with 1% BSA ASW. They were washed in PBS (1.2 g Tris, 6 g NaCl, 0.2 g KCl/l, pH 7.4) several times, 35% glycerol solution was added, and then observed as described above. Twenty embryos were measured using a micrometer each hour from one batch of *T. toreumaticus*, two batches of *T. reevesii* and *M. globulus*, and four batches of *T. hardwickii* according to Kominami and Masui (1996) (Fig. 1). Each embryo’s total length and width was measured, the total length of the archenteron, the diameter of the blastopore of the archenteron, the outer or inner diameter of the archenteron at the middle part of the total length of the archenteron and the outer or inner diameter of the archenteron on the tip (In *T. hardwickii*, embryos 7 h after initiation of invagination were not measured).

**Table 2.** Summary of gastrulation of four temnoleurids.

| Species | Factors of elongating archenteron |  |  |  |  |
| --- | --- | --- | --- | --- | --- |
|  | Invagination<br>type | Cell<br>migration | Cell<br>elongation | Cell<br>rearrangement | Towing by<br>SMCs |
| *T. toreumaticus* | Continuous* | Yes | Yes | Yes | No? |
| *T. reevesii* | Stepwise | No? | No | Yes | Yes? |
| *T. hardwickii* | Stepwise | Yes | No | Yes | Yes? |
| *M. globulus* | Stepwise* | Yes | Yes | Yes | No*/Yes? |

\*The report of Takata and Kominami (2004) also showed that *T. toreumaticus* invaginates continuously and *M. globulus* invaginates stepwise without conspicuous rearrangement.

## Acknowledgements.

We appreciate the Department of Fishery in Yamaguchi Prefecture and Yamaguchi Fisheries Cooperative Association for permission to collect sea urchins.

## Author contributions

C. K. and A. Y. designed and T. F. and Y. E. performed the experiments equally. C. K. and A. Y. wrote the manuscript by discussion with M. K.

